# REST: A method for restoring signals and revealing individual macromolecule states in cryo-ET

**DOI:** 10.1101/2022.07.11.499538

**Authors:** Haonan Zhang, Yan Li, Yanan Liu, Dongyu Li, Lin Wang, Kai Song, Keyan Bao, Ping Zhu

## Abstract

Cryo-electron tomography (cryo-ET) is widely used to explore the 3D density of biomacromolecules. However, the heavy noise and missing wedge effect prevent directly visualizing and analyzing the 3D reconstructions. Here, we introduced REST, a deep learning strategy-based method to establish the relationship between low-quality and high-quality density and transfer this knowledge to restore signals in cryo-ET. Experimental results on purified ribosome and recombinant nucleosome datasets showed that REST had outstanding performance in denoising and compensating the missing wedge. The application in dynamic nucleosome structures suggests that REST has the capability to reveal individual macromolecules which present different conformations without subtomogram averaging. Moreover, REST could greatly improve the reliability of particle picking. These advantages enable REST to be a powerful tool for the straightforward interpretation of target macromolecules by visual inspection of the density and of a broad range of other applications in cryo-ET, such as segmentation, particle picking, and subtomogram averaging.

## Introduction

Cryo-ET has emerged as a powerful method which could record the 3D information of the biological macromolecules; however, many challenges still remain to be addressed (Beck and Baumeister, 2016; Li, 2021). First, the noise level of the tomogram is very high due to the high radiation sensitivity of the samples; hence low-dose electron tomography, which hinders human eyes to identify the features in it (Hattne et al., 2018). Second, during the data collection, tilt-series images can only be collected within a tilt angular range of approximately ±70° because of the limitation of the specimen holder. This could lead to incomplete 3D information in the Fourier space, resulting in a so-called missing wedge in the tomogram. The effect of the missing wedge is clearly visible in the 3D Fourier transform of the beam direction. The most obvious artefact caused by a missing wedge is the anisotropic resolution, which means that objects appear elongated in the direction of the beam axis, i.e., in the Z direction (Moebel and Kervrann, 2020). The EM density in the 3D and 2D slices related to the Z-plane are distorted as a result of this elongation. Therefore, most of 3D segmentation was unable to entail in Z direction and render a highlight extended structure.

To address these challenges in cryo-ET, a variety of methods have been proposed to recover the information and produce high SNR tomograms (Sorzano et al., 2017). During the data collection, dual-axis tomography, in which the tilt series is collected using two perpendicular axes, could be applied (Mastronarde, 1997). However, this method is limited by the use of a higher electron dose, which may damage the biological specimen (Guesdon et al., 2013). In other studies that have focused on the data processing procedures, a series of algorithms, including the algebraic reconstruction technique (ART) (Gordon et al., 1970), simultaneous ART (SART) (Andersen and Kak, 1984) and simultaneous iterative reconstruction technique (SIRT) (Agulleiro and Fernandez, 2011), have been proposed to improve the quality of tomograms. These methods, which are mainly based on mathematic calculations, reduce the differences between the calculated projections of the reconstructed tomogram and the tilt series. By using these algorithms, high contrast for visualizing 3D structures can often be achieved from the tomogram in which the low-resolution information is retained while high-resolution information is discarded. In addition to the above algorithms, the compressed sensing (CS)-based method has also been proven to be effective in recovering the information in electron tomograms (Böhning et al., 2022; Deng et al., 2016; Leary et al., 2013). It introduces *a priori* assumptions in the tomogram, e.g., density positivity and solvent flatness, to constrain the structural features and allow the high-fidelity reconstruction of signals. By applying CS on biological samples, ICON was found to be capable of reconstructing tomograms with high contrast and successfully restoring the missing information (Deng et al., 2016). A more recently proposed method, CS-TV^2^, which uses an advanced CS algorithm, could increase the contrast while retaining high-resolution information (Böhning et al., 2022). However, CS-based methods rely heavily on sufficient signal-to-noise ratio (SNR) and thus require high-contrast tomograms.

In recent years, deep learning algorithms have been increasingly applied in cryo-EM and cryo-ET workflows (Bepler et al., 2019; Wagner et al., 2019). Learning-based methods, e.g., Topaz-Denoise (Bepler et al., 2020), have been shown to be advantageous in denoising tomograms. It presents a general 3D denoising model of noise2noise (N2N) for improving tomogram interpretability. In addition to denoising tomograms, deep learning has also been used to recover missing-wedge information. In a recent study, a joint model that was designed to recover the missing-wedge sinogram was proposed (Ding et al., 2019). It required a U-Net structure combined with a generative adversarial network (GAN) to reduce the residual artefacts. However, the proposed joint model was still limited to 2D data due to the lack of ground truth for the training network model in cryo-ET. In addition to the aforementioned joint model, an application named IsoNet, which learns from the information scattered in the original tomograms with recurring shapes of molecules based the U-Net framework, is used to recover missing wedges in cryo-ET (Liu et al., 2021). However, as the processing of IsoNet, its “ground truth” still preserves noticeable artefacts with a high noise level. Since the training does not employ the real ground truth of each density directly, the effect of restoring relies on the feature and SNR in the density mask. Actually, the raw tomogram suffers from noise and missing wedge which are irreversible, thus, it makes the acquisition of ground truth very challenging. Therefore, for deep learning strategies aiming at information restoration, it is critical to generate suitable training datasets to train the neural network.

In this study, motivated by the joint model and IsoNet, we proposed a knowledge transfer method for **<u>re</u>**storing the **<u>s</u>**ignal in **<u>t</u>**omograms (REST) to denoise the tomograms and compensate for the missing-wedge information in cryo-ET. To address the issue of the nonexistent ground truth to train the neural network, training pairs were generated from two strategies depending on the different specimen homogeneity. The network model after training was found to be capable of restoring a sparse and elongation-free volume from the noisy tomograms with severe artefacts. As REST is highly robust to noise, the raw tomogram generated from weighted back projection (WBP) can be directly taken as the input, and preprocessing such as deconvolution or filtering is not required to improve the SNR. By applying the REST method to purified ribosome and recombinant nucleosome datasets, we found that it is highly capable of denoising the tomograms and compensating for the missing-wedge information. With a significantly improved view and the knowledge of conformational changes, REST helps us to identify the target macromolecules with different conformations in both simulated datasets and real specimens. The results show that REST can greatly enhance the visualization of macromolecules and improve the structural interpretability of cryo-ET without subtomogram averaging (STA).

## Results

### Workflow of REST

We use U-Net modified from IsoNet, from which the relationship between the input volume (low-quality density) and the ground truth (high-quality density) can be learned, as a model for segmenting dense volume from sparse annotation. The general workflow of REST is comprised of three parts, i.e., generating training pairs, training the model and restoring information, as depicted in Fig. 1.

**Fig. 1.**
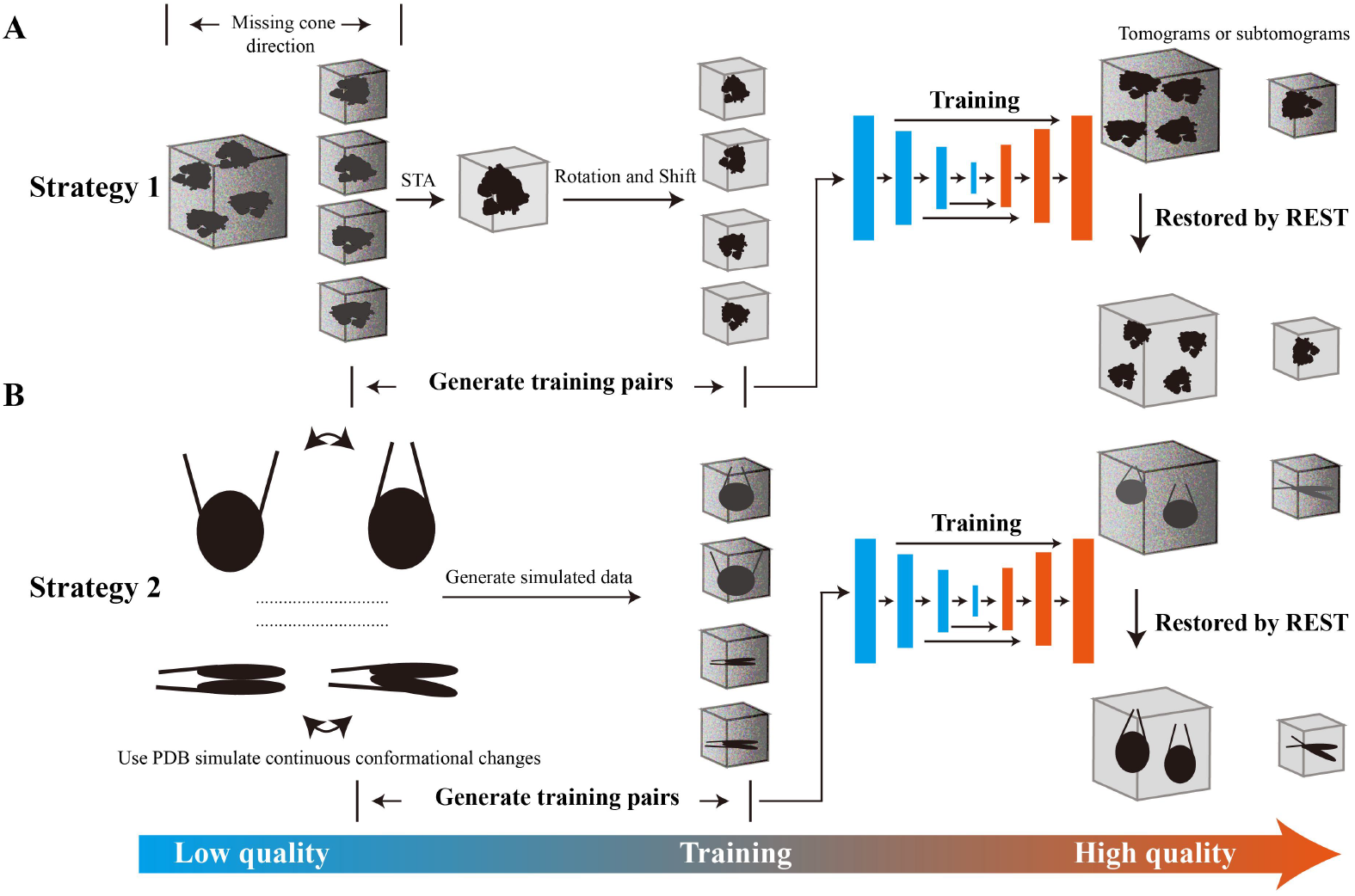
The workflow of the REST method for restoring signal. **(A)** Strategy 1 consists of generating the training pairs using subtomogram averaging (STA), training the model and restoring information. **(B)** Strategy 2 consists of generating the training pairs using simulated data, training the model and restoring information. The noise volume images appear distorted. The clean volume images have no distortion with high SNR.

### Generating training pairs

Two strategies were proposed to generate training pairs.

Strategy 1: Subtomogram averaging based strategy

Step 1: Subtomogram averaging

Subtomogram averaging was performed by using limited amounts of particles for missing-wedge compensation and elongation-free map generation with a high SNR. The accurate alignment parameters for each particle could be used to efficiently reduce the loss in training. The averaged map was used as the ground truth for each particle.

Step 2: Extraction of subtomograms

The subtomograms that were averaged in step 1 were extracted as the input of the training data.

Step 3: Generation of the ground truth

According to the alignment parameters from each subtomogram, the averaged map was rotated and shifted to generate the ground truth. By coupling the input of training data from step 2 and the ground truth, the training pairs were obtained.

Strategy 2: Simulation-based strategy

Step 1: Generation of dynamic models using normal mode analysis (NMA)

Normal mode analysis (NMA), a method for molecular mechanics simulation (Tama et al., 2000), was used to generate dynamic models from a static (pseudo) atomic model. Based on the prior knowledge of the target molecule, the conformations of interest were selected among these dynamic models.

Step 2: Generation of the ground truth

We converted the selected dynamic models to the EM density and then rotated and shifted the density in the 3D space using random Euler angles and shifts as the ground truth.

Step 3: Generation of the simulated data

The tilt series were obtained from the EM density of step 2. By adjusting the SNR and modulating the images with the contrast transfer function (CTF), the volume was reconstructed and used as the simulated data. It was also used as the input of the training data. Details of the method used to simulate the data are provided in the Methods section. By coupling the input of the training data and the ground truth from step 2, the training pairs were obtained. Usually, both strategies 1 and 2 can be applied to the stable target objects. However, strategy 2 is more suitable for the target object with noticeable dynamics.

### Training the model

We employed a U-Net-based voxel-wise network derived from IsoNet. One of the advantages of U-Net is its ability to segment the dense volume from sparse annotation (Çiçek et al., 2016). Therefore, U-Net is particularly suitable for segmenting parse features from cryo-electron tomograms containing heavy noise and elongated artefacts due to the missing wedge effect. The main blocks in U-Net are built from stacking multiple layers, which are used for 3D convolution and deconvolution. The convolution and deconvolution layers are used for extracting the features of target objects and recovering the high-resolution features. After training, the mapping relationship is established between the low-quality particles and the ground truth (i.e., the high-quality particles) with their corresponding orientations. The mapping relationship and knowledge learned from the training pairs can then be transferred to restore the low-quality real density.

### Restoring information

To evaluate the robustness of REST when it is influenced by missing information and noise, we tested the restoring capability of REST using a series of data (PSIM01, 02, 03, 04), which were simulated under different conditions. Interestingly, we found that REST could handle the disturbances of noise and missing wedges well, and good performance was achieved in restoring information even when the SNR was reduced to 0.01 and only a tilting range of −40°~ 40° was implemented. The correlation coefficient (CC) between the prediction and the ground truth was close to 1.0 (Fig. S1), suggesting an almost complete restoration from the noisy volume in the predicted particle. Notably, the input volume for restoration does not require preprocessing steps such as deconvolution or filtering to improve the SNR, which means the raw reconstructed tomogram from WBP could be selected as the input. Using the raw WBP reconstructed tomograms as the input has an advantage when retaining the high-resolution features, as most of the preprocessing or denoising steps in tomograms, such as low-pass filtering, removes these high-resolution signals and results in an adverse influence on restoration.

### REST shows the capability to enhance SNR and reduce resolution anisotropy in real data

To evaluate the restoration capability of REST on the (sub)tomograms after training, we first applied it to the EMPIAR-10045 dataset, which contains the tomograms of ribosomes (~ 25nm), stable samples with high abundances *in vivo*. For this dataset, strategy 1, i.e., the subtomogram averaging (STA)-based strategy, was used to generate the training pairs. We directly extracted the particles from the raw tomograms as input, calculated an STA averaged map and rotated and shifted the average map corresponding to the orientation of each particle as the ground truth (Fig. 2 A). After training the model and restoring information using REST, we found that the corresponding missing-wedge information was significantly recovered in the Fourier space (Fig. 2 B) and the restored tomograms (EM01) were highly visible (Fig. 2 C).

**Fig. 2.**
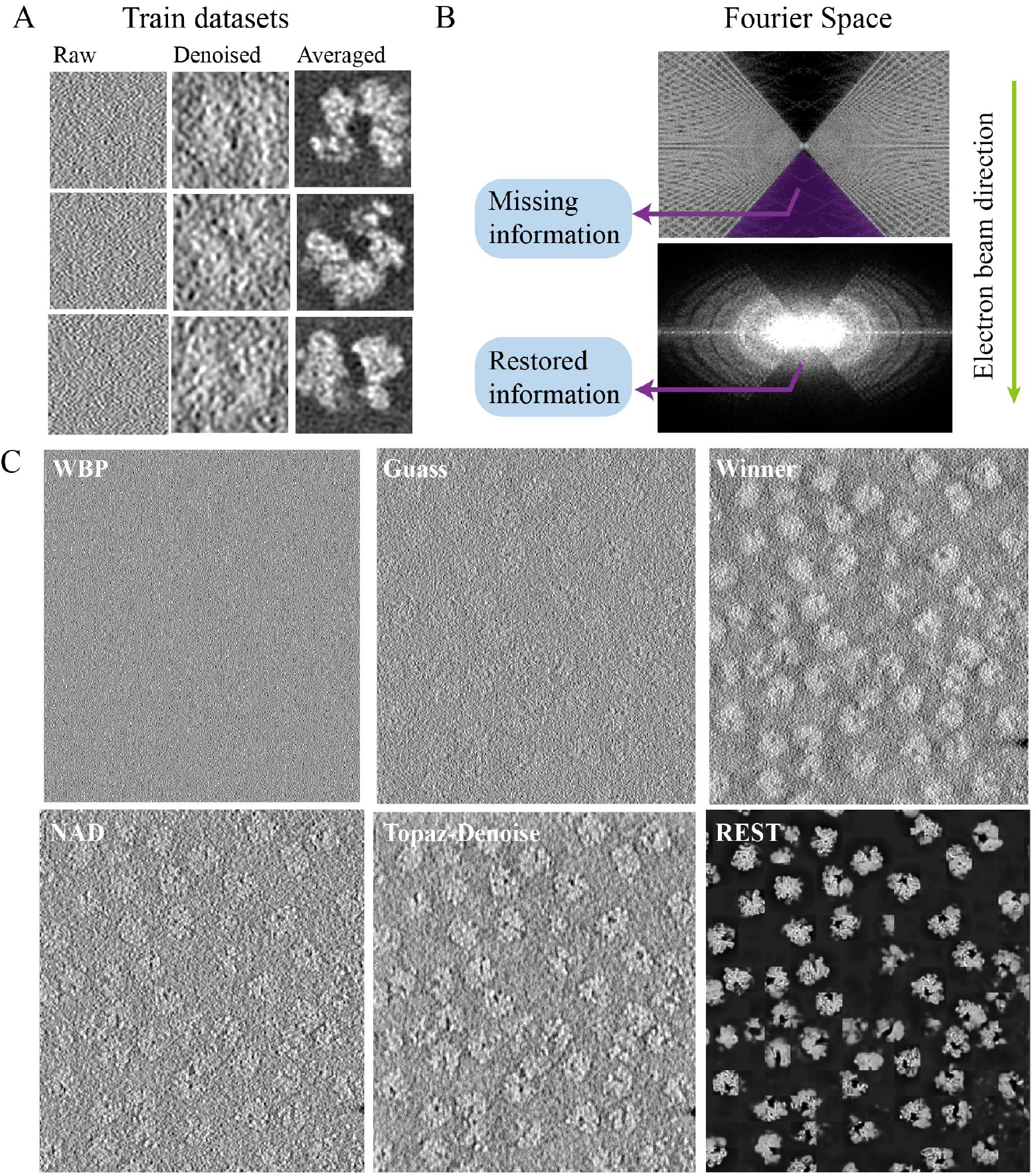
Results from the real datasets of ribosomes (EMPIAR-10045) using REST. **(A)** Examples of three training pairs. Left: the raw particles extracted from the tomograms (input); Middle: the coarsely denoised particles, which are easier to observe; Right: the averaged map that rotated and shifted based on the alignment parameters. **(B)** Fourier transforms of the raw data and restored tomogram. **(C)** Comparison of the raw data (WBP) and the denoised data using the Gaussian filter, Winner filter, NAD, Topaz-Denoise and REST methods. The REST method could thoroughly remove noise that appears in the tomographic slices.

Compared with other denoising methods, such as the Gaussian filter, Winner filter, Topaz-Denoise, nonlinear anisotropic diffusion (NAD), we found that REST achieved a stronger noise removal performance and reserved more signals in the 2D slices (Fig. 2C). We quantitatively assessed the denoising performance by measuring the SNR of raw slices and slices denoised with these methods. Since the ground truth was not available for the datasets, the SNR was estimated in a similar approached that in Topaz-Denoise (Bepler et al., 2020). First, we averaged ten slices into one micrograph. Then, we selected 10 paired signal and background regions across the micrographs. Given the signal N and background pairs 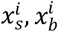, the mean and variance of each background region is 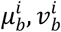. We defined the signal for each region as 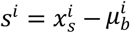 and calculated the mean and variance of the signal region, 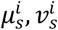. The average SNR in dB for the regions is defined as:

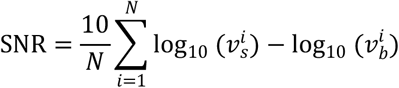

As shown in Table 1, the SNR was improved by approximately 0.5 dB over the raw micrographs when using the conventional methods. Notably, the SNR was improved by 7 dB over the raw slices and approximately 6 dB over other methods when using REST, which indicates that a significant improvement in SNR enhancement is achieved.

**Table 1:** Comparison of denoising methods based on the estimated SNR (in dB, larger is better)

| SNR | Dataset |  |
| --- | --- | --- |
|  | EM01 (ribosome) | EM02 (nucleosome) |
| *Method* |  |  |
| Raw data | 0.352 | 0.05 |
| Gauss filter | 0.419 | — |
| Winner filter | 0.598 | — |
| NAD | 0.652 | 0.14 |
| Topaz-Denoise | 0.815 | 0.23 |
| REST | 7.337 | 9.57 |

Additionally, after restoration by REST, we found that each particle could be identified clearly not only in the XY-plane but also in the XZ-plane, with few elongation artefacts (Fig. 3A). Thus, REST enables the accurate identification of particles in all directions. For the nonribosome densities, such as those of carbon films, the false-positive densities were not segments by REST (Fig. 3A). In addition to the 2D slices, compared with the density processed by Wiener filtering (in order to visualize the density), REST was able to restore the 3D density of the particle (e.g., the green and yellow particles) with almost no visible elongation and distortion (Fig. 3B).

**Fig. 3.**
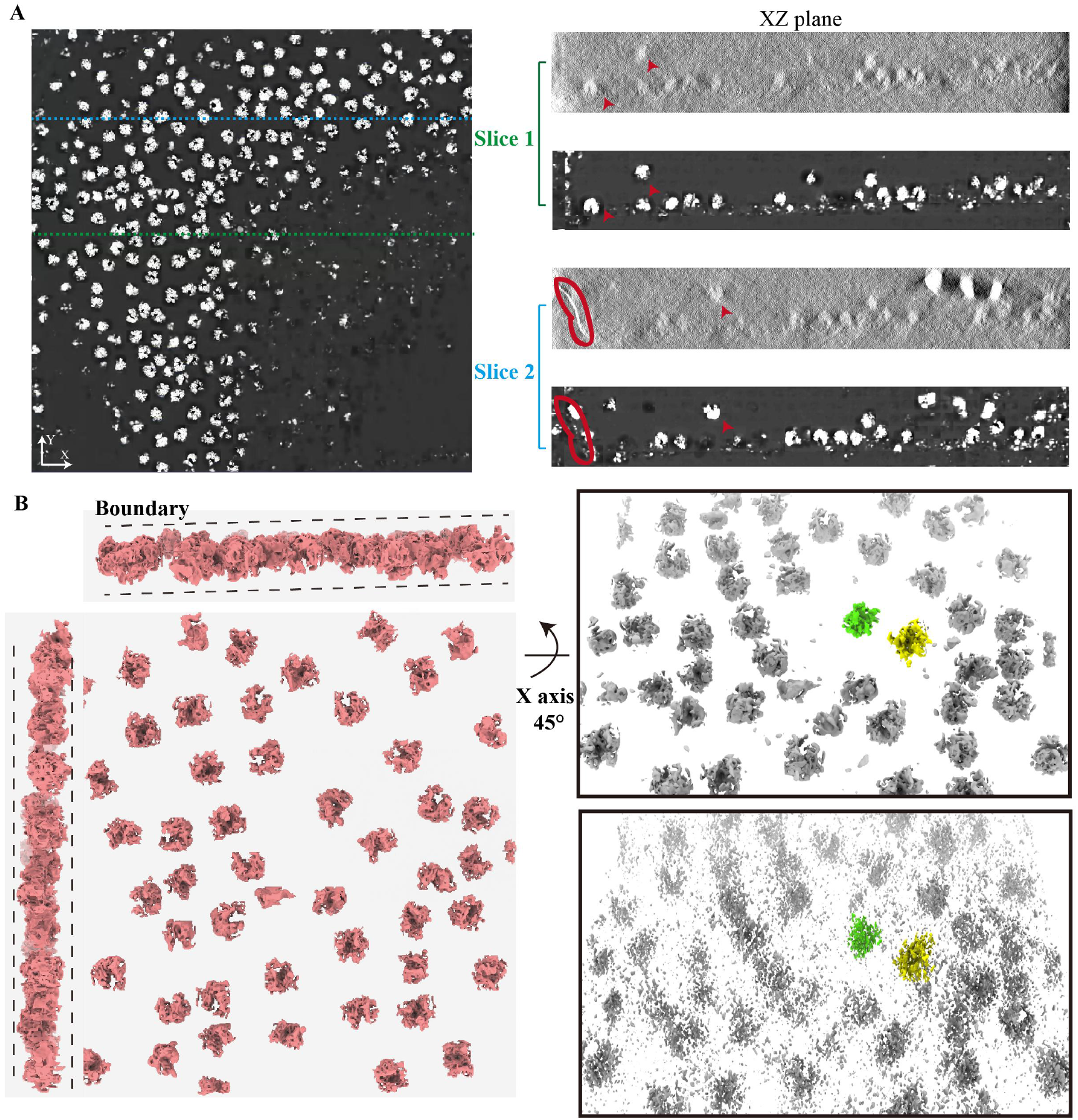
REST enabled the restoration of 3D density and the accurate identification of particles in all directions. **(A)** Left panel: Views of the XY-slice of the tomogram restored by REST; Right panel: Two XZ-slice views of conventionally denoised (Winner filter) tomogram (top) and REST restored tomogram (bottom). The red arrows indicated the corresponding ribosomes. The red ellipse indicated the edge of the carbon film, and there is no corresponding false-positive density in the restored tomogram. **(B)** Left panel: A 3D rendering of the tomogram restored by REST. Right panel: The tomogram restored by REST (top) had a better restoration capability than that in the conventionally denoised tomogram (Wiener filtering) (bottom). Both the tomograms were rotated around the x-axis by 45° corresponding to left panel.

In addition to strategy 1, strategy 2, i.e., the simulation-based strategy, was also tested to train the network. This strategy was applied to the EM02 dataset of nucleosomes (~ 10 nm) that we reconstituted. The reason we selected nucleosomes is that nucleosomes have area notably smaller size than the ribosomes studied above, and are known to be highly dynamic (see below). A similar improvement in terms of compensating for the missing wedge and denoising the tomogram was noted on the nucleosome dataset (Table 1, Fig S2). These results suggest that both strategy 1 and strategy 2 could be implemented to achieve an enhanced SNR and reduced resolution anisotropy and could be successfully applied to real tomograms.

### Application of REST to simulated flexible sample revealed conformational changes in the individual particles

It is known that most macromolecules are not strictly rigid but are flexible entities with continuous conformational transitions when performing their biological functions. Although the STA method can be used for classification to study different conformations, particles with continuous conformations in the subtomograms are rarely assigned to the same class. In addition, the number of particles in each class is insufficient to obtain a high-quality averaged result. Thus, the complicated cell environment and continuous conformational changes of the specimen make disentangling the data heterogeneity by STA difficult. Here, by using NMA, we generated a simulated dataset of 177 bp nucleosomes that have flexible flanking linker DNA and continuous conformational changes as a test object (Fig. 4A). The training pairs were obtained using strategy 2 as shown in Fig. 4B. Notably, after training, the test densities restored by REST were highly consistent with those of the ground truth (Fig. 4C).

**Fig. 4.**
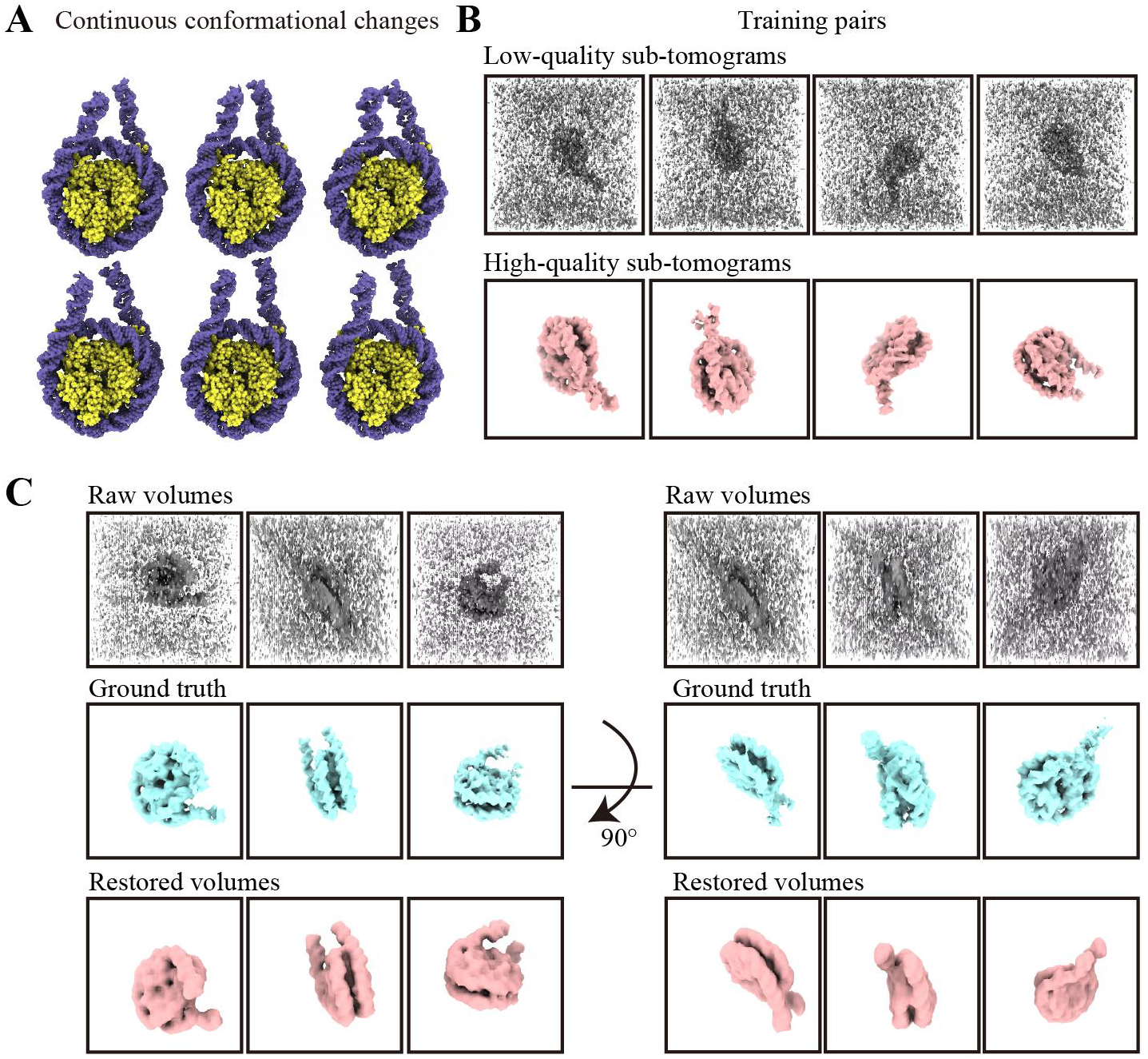
REST revealed continuous conformational changes of nucleosome linker DNA in simulated dataset. **(A)** A series of dynamic nucleosomes generated by using NMA. **(B)** Examples of four training pairs. Top: the input of training pairs generated from the corresponding ground truth particle with noise (SNR 0.1) and a missing wedge (±40°) superimposed (low-quality). Bottom: the ground truth of the training pairs that are generated from the atomic model (high-quality). **(C)** The 3D volume of input particles (raw volumes) (top), the corresponding ground truth (middle) and the restored volumes using REST (bottom).

Remarkably, we found that REST could be also used to discern a series of conformational changes in the tomogram. For example, in the TSIM01 dataset, the REST-restored tomogram was highly consistent with the ground truth, and the contrast was significantly improved compared with the raw data. In the Fourier space, the missing wedge is also compensated well (Fig. S3). In addition to the 2D slices, REST also restored the 3D density, and the structural variation, e.g., the linker DNA breathing motions between the open and closed states (Zhou et al., 2019), could be clearly visualized and unambiguously identified (Fig. 5A and B). By analysing the missing-wedge information, we found that REST can effectively eliminate elongation and distortion, which shows a significant improvement (Fig. 5C). These results suggested that REST could be used to directly identify and display the conformational changes of the dynamic structures.

**Fig. 5.**
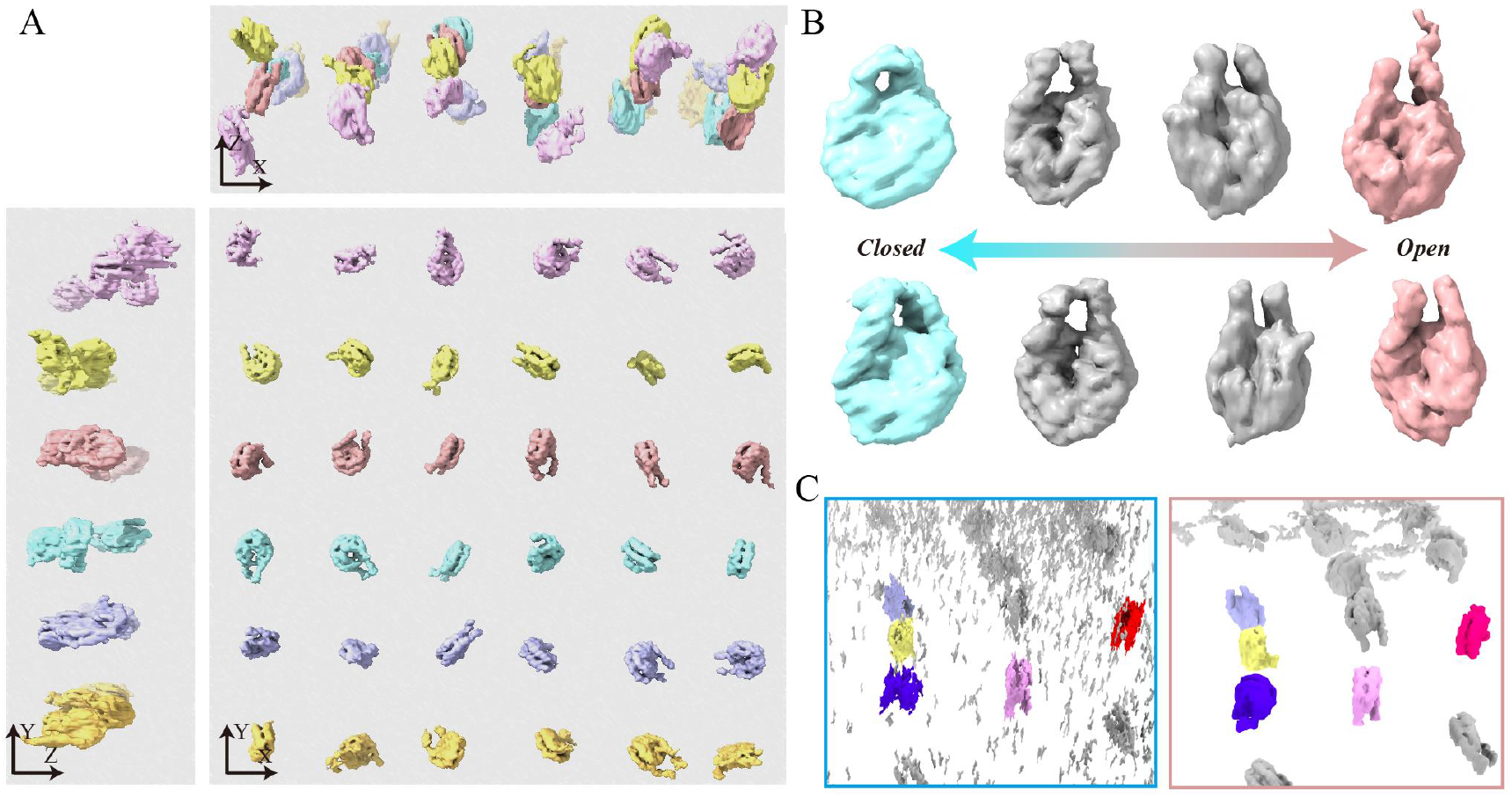
REST identified and displayed the conformational changes of nucleosome linker DNA in simulated data. **(A)** A 3D rendering of the restored tomogram. Each nucleosome could be identified. **(B)** Eight representative particles from (A) showing the motion of linker DNA from closed state (cyan) to open state (pink) *via* middle transitions (gray). **(C)** Compared with the 3D density improved by Topaz-Denoise and IsoNet (left), REST could effectively clean the noise density and eliminate elongation and distortion (right). The region was from tomogram in (A) and rotated around the x-axis by 45°.

Interestingly, we found when using REST, the data for training did not necessarily include all the possible conformations of the particles. In this study, we only used limited amounts of conformations to train the network. Nevertheless, many conformations that were not included in the training dataset could still be identified. This result indicates that REST could transfer knowledge from limited prior information to analogous information of a broad cast.

### Applying REST to real nucleosome data with different lengths of DNA reveals the individual characteristics of the particles

In addition to the simulated nucleosome dataset, real tomograms of nucleosome samples with linker DNA (the EM03 dataset) were also tested. We reconstituted nucleosome with linker DNA particles, mixed them with nucleosome core particles (NCP) without linker DNA as a control, and collected a series of electron tomography datasets for testing REST. According to the result, we found REST also presented a notable improvement in SNR and recovering the missing wedge in Fourier space (Fig 6A). The restored tomogram was shown in Fig. S4. We used the combination of Topaz-Denoise and IsoNet to denoise and compensate for the missing wedge in tomogram, which is referred to as the T-I density hereafter. Compared with T-I density, the 3D density of each subtomogram after REST restoration was less elongated and distorted, and thus, closer to the real structure (Fig. 6B). We also statistically analysed the CC value between the density restored by REST and the T-I density (Table S2). The high CC value achieved by REST indicated that the restored densities were derived from the raw tomogram. As a consequence, nucleosomes with linker DNA could be readily distinguished from NCP by visualizing the flanking linker DNA out of the nucleosome (Fig. 6B). Compared with the wrapped DNA on the NCP, the extra unwrapped linker DNA was apparently flexible, and thus, showed versatile conformations. By applying REST to the nucleosomes with flanking linker DNA, we could distinguish the symmetric and asymmetric linker DNAs with extended or curved conformations that coexist within the nucleosomes (Fig. 6B). This kind of structural flexibility is consistent with the nucleosome variations in interphase and metaphase chromosomes (Arimura et al., 2021). These results indicated that interpretable information, such as dynamically changing nucleosomes with different conformations, could be directly derived from the elongated and noisy subtomogram after restoration by REST.

**Fig. 6.**
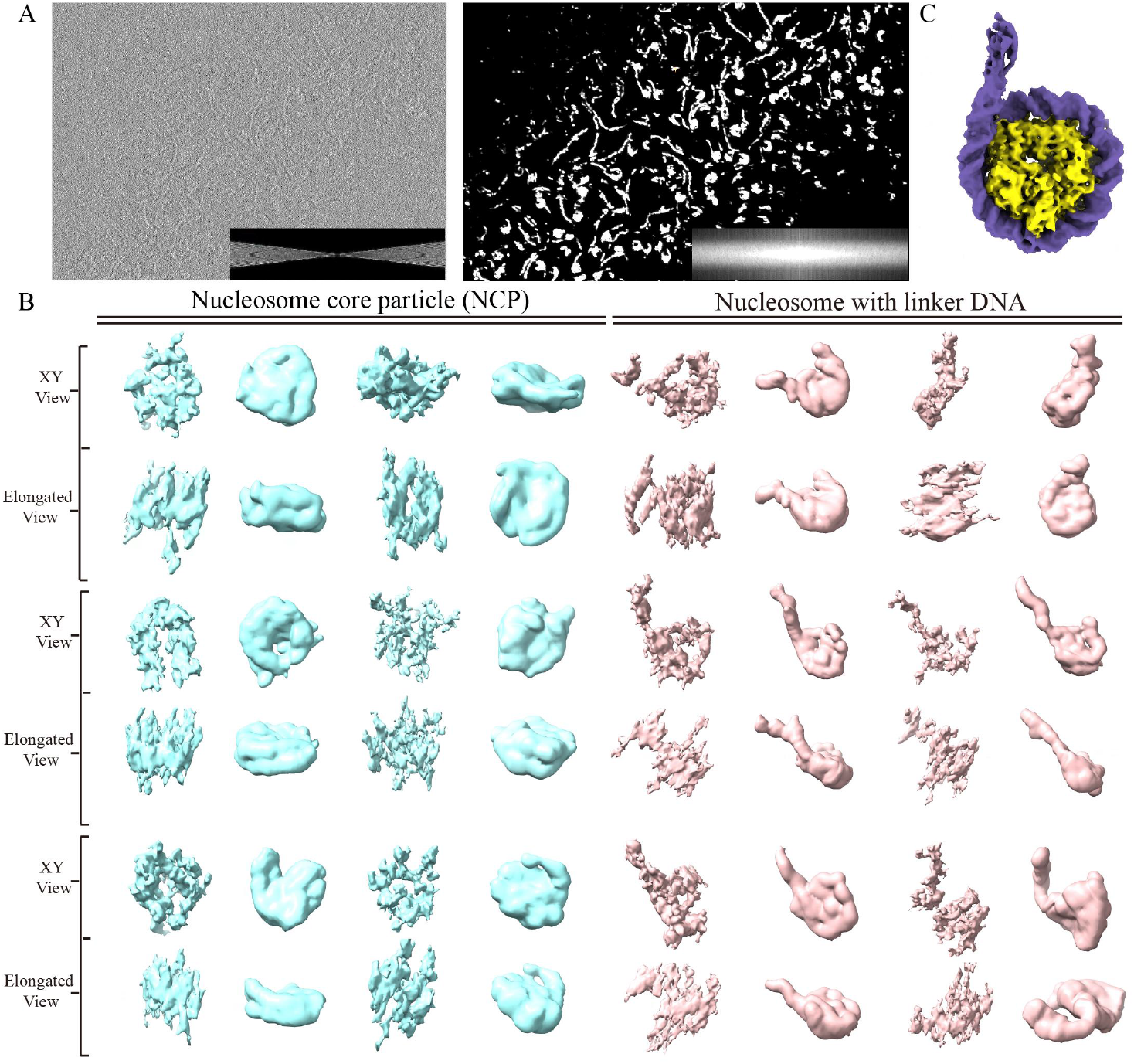
REST could directly reveal versatile characteristics of dynamic nucleosomes and free DNA. **(A)** Left: The 2D slices of the raw tomogram; Right: REST-restored tomogram. The Fourier transforms are shown in the right corners. **(B)** Comparison of volumes denoised by the combination of Topaz-Denoise and IsoNet (T-I density, left) and REST-restored density (right). For clarity, nucleosome core particle and nucleosomes with linker DNA contains two different views: the XY-view and the tilted view with the elongation (elongated view). **(C)** The 3D map of the reconstituted nucleosome with linker DNA by conventional single-particle reconstruction shows the asymmetric linker DNA.

In addition to the REST method, we also applied conventional single particle analysis (SPA) to reveal the 3D structure of reconstituted nucleosomes with linker DNA. Subjecting approximately 200,000 particles to averaging, the structure was resolved at a resolution of 3.7 Å, which showed an asymmetric linker DNA at one end of the nucleosome (Fig. 6C). Interestingly, this specific conformation, as shown in the structure resolved at high resolution by SPA, was also found in the architectures discerned by REST without averaging. Therefore, the results indicated that different from the conventional averaging method which using large amounts of particles and mainly revealed an averaged character of particles, the REST method presented in this study could directly reveal versatile characteristics of target macromolecules, thus had a great practical application in cryo-ET.

### Restoration by REST facilitated particle picking in cryo-ET

To free researchers from particle picking work on tomograms, a number of methods have been proposed (Moebel et al., 2021; Wagner et al., 2019). Usually, the template matching method is the first choice if a template is available. However, this method suffers from missing wedges and noise; thus, the calculated CC value between the subtomogram and template is relatively low. Consequently, false-positive hits and unreliable results often occur. Since the REST method can be used to achieve both enhanced SNR and missing wedge compensation, it can also be used as a preprocessing method in particle picking before template matching. We tested both simulated data and the corresponding REST-restored data for template matching. The statistical offset from the ground truth centre and the CC value between the subtomogram and template were compared to evaluate the performance of REST restoration on picking particles.

As shown in Fig. 7A, the coordinates calculated from the tomograms restored by REST are extremely consistent with the ground truth centre. In contrast, the coordinates calculated directly from the raw tomogram present variant deviations, although most centres of particles are identified right. However, as shown in Fig. 7B, the CC values calculated from the two tomograms are noticeably different. The CC value, which reflects the confidence of the particle centre and the orientation, was significantly improved in the tomogram restored by REST. This was most likely contributed to the high consistency between the restored density and the real signal. These results indicate that using tomograms restored by REST could greatly improve the reliability of particle picking with a very high CC value.

**Fig. 7.**
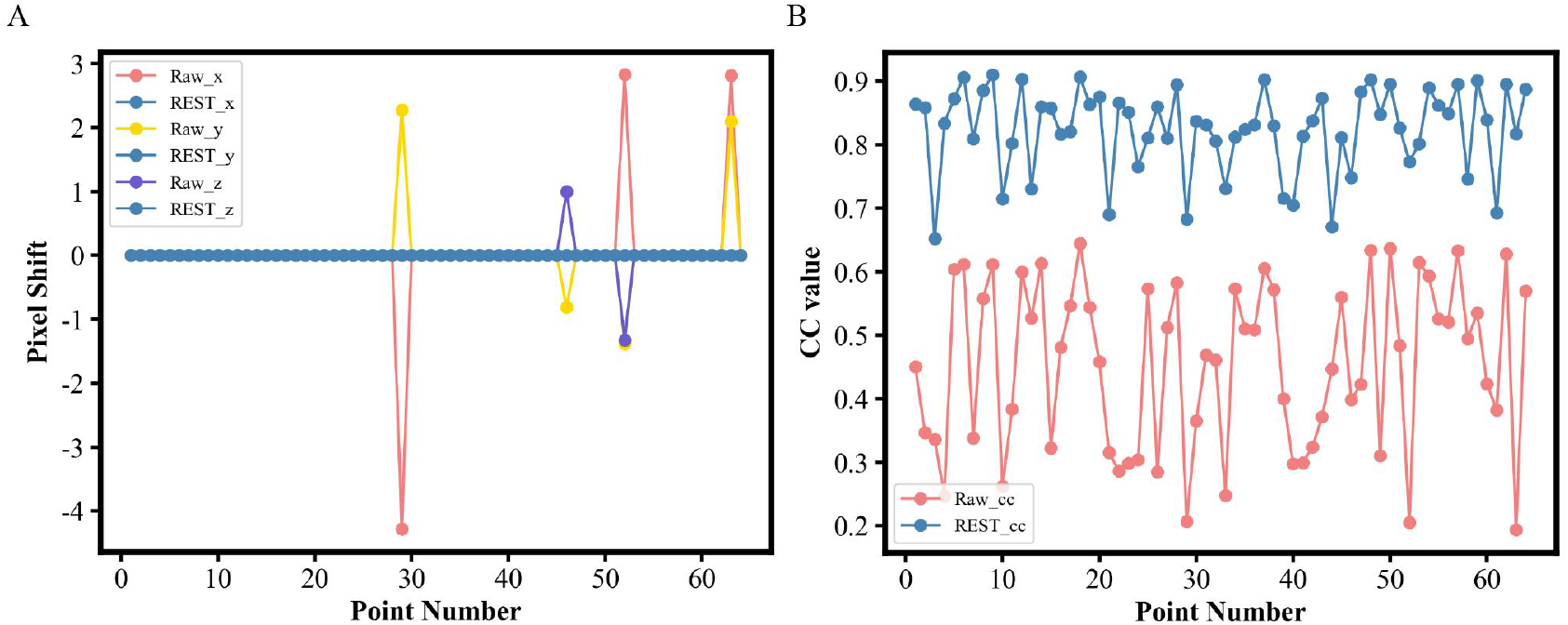
Comparison of particle picking reliability between the raw tomogram and the REST-restored tomogram. **(A)** The offset of each picked particle shown in the X-, Y-, and Z-directions in the raw tomogram and REST-restored tomogram. **(B)** Comparison of the CC values calculated between the picked particles and templates in the raw tomograms and tomograms restored by REST.

## Discussion

Cryo-ET has been increasingly used in 3D structural studies of native biological samples (Erdmann et al., 2021; Obr et al., 2022; Wang et al., 2022). By reading reconstructed tomograms, biomacromolecule information, including its spatial arrangement, architecture, or even specific orientation, is expected. However, due to both the low SNR and missing wedge effect, the interpretation of the tomogram is largely limited. Sometimes even the identification or classification of the target biomacromolecule is notoriously difficult. To overcome this drawback, subtomogram averaging, which is laborious, and challenging, has often been necessary. In addition, this kind of averaging method usually requires disentangling the continuous architecture from the flexible samples. In contrast, the REST method presented in this study could be used to significantly enhance the SNR and reduce resolution anisotropy without averaging. This could greatly help to reveal the individual characteristics of each biomacromolecule. The rendered architecture also produces a clean density that is sufficient enough for us to distinguish fundamental features, thus making the (sub)tomogram directly interpretable.

It is well known that deep learning relies heavily on training datasets. Thus, using real data with the ground truth for training is the best choice to ensure the network performance. However, the raw tomogram suffers from noise and an irreversible missing wedge, making the acquisition of the ground truth very challenging. STA provides an alternative approach to obtain the ground truth of the raw data. This approach could be used to consequently establish a mapping relationship between the raw data and an averaged map for restoration. Nevertheless, as most macromolecules are flexible, the averaged structures present insufficient features, and thus, a valid mapping relationship is rarely established. To address this situation, in this study, we found an alternative strategy that conversely degrades high-quality data to simulate low-quality data that are close to the raw data and establish a relationship between them. As long as the neural network can learn the mapping relationship well, the model can be migrated into raw data to generate the corresponding high-quality data. Apparently, the most challenging issue is how to make the simulated data closely related to the raw data. We found that the analogous contrast and elongation are the keys to simulating subtomograms in cryo-ET. Applying the conformational changes in datasets would also learn more knowledge of flexibility and greatly improve the restoration ability.

It is worth noting that REST uses the U-Net framework, which is widely used in segmentation. In most of segmentation methods, e.g., EMAN (Chen et al., 2017), the features are manually or automatically labelled in 2D slices. In the REST method, the receptive field is boarded to 3D. The perceptron synchronously globally to locally learns the relationship of the two maps in 3D; thus, the model could eliminate the artefact found in 3D. Moreover, most of the methods for segmentation are highly sensitive to the SNR. Interestingly, in REST, the preprocessing of the input dataset is not needed, i.e. the raw tomograms from WBP could be directly used for restoring.

As mentioned above, REST can be used to reveal each particle state without the need for time-consuming STA. However, it is worth noting that REST cannot retain the high-resolution information from raw data. This is because the process of training which is essentially a regression problem needs to reduce the error between the ground truth and restored volume. In practice, the low-frequency information accounts for the majority of the signal, whereas the high-frequency signal is under the noise and hardly to label accurately. Incorrect results will backproject and update the weight in network. This process leads the loss of high-frequency information in order to convergence the loss function. Nevertheless, in many situations, the restored density is sufficient for the distinction of the shape and other fundamental features, as shown in the above examples.

In conclusion, REST presented in this study provides a way to enable the direct observation of fundamental architectures and conformational changes for functional interpretation without the laborious and challenging averaging process. Thus, it could be of broad utility to the cryo-ET community by the function of restoring a clear signal like picking particles in a noisy background, segmenting the target feature, identifying dynamic or flexible architectures, obtaining the density without elongation as the initial reference for STA and even guiding the particles to be classified and aligned for STA.

## Materials and methods

### Implementation specifics of the REST method

In strategy 1, to process the ribosome data used in this study, the STA steps were followed according to the protocols in Relion (Bharat and Scheres, 2016). Two tomograms in EMPAIR-10045 were used to perform STA, and the other volumes were used as the test data. The averaged subtomograms were first extracted using Relion (Zivanov et al., 2018) and then used as the input to the training data. According to alignment parameters in star files, *e2proc3d*.*py* (Tang et al., 2007) was used to rotate and shift the averaged map to generate the ground truth. By combining the raw particles and the averaged maps, which were reset to the corresponding orientation, the training pairs were obtained.

In strategy 2, NMA was implemented in the target object. According to the prior of the target model, a series of atomic models were generated by NMA, in which possible conformations were selected. The atomic model was converted to the EM density using *e2pdb2mrc*.*py* (Tang et al., 2007), and the density was then rotated and shifted in the 3D space using random Euler angles and random x, y, z shifts as the ground truth. The simulated data and training pairs were then generated.

### Simulating datasets based on experimental parameters

To generate the simulated dataset for each subtomogram or tomogram, the following steps were performed:

a. Rotate the obtained averaged map or the density from the atom model at random Euler angles, with random x, y, and z displacements (*e2proc3d*.*py*). For example, by using the density converted from the atom model, 3000 subtomograms (64 voxels, 4.4 angstroms) were generated.
b. Project the ground truth according to the corresponding collection conditions (e.g., ±60°, 2°) by using the *relion_project* toolbox.
c. Perform CTF modulation on each projection. Gaussian noise is added by using *xmipp_phantom_simulate_microscope*, and then the CTF phase is inverted.
d. Reconstruct the tilt series using *relion_reconstruct* to obtain the simulated data with missing wedges and noise.

## Datasets

Six simulated datasets and three real datasets were used to evaluate the performance of REST. We generated one simulated dataset by using the averaged map of the ribosome as PSIM01, four simulated datasets by using pdb:3AFA as PSIM02-05 and one simulated tomogram by an atom model of 177 bp nucleosome as TSIM01. Note that PSIM stands for simulated subtomograms and TSIM stands for simulated tomograms. We downloaded EMPIAR-10045 (the purified ribosome dataset) as EM01, collected the tomogram of the 147 bp recombinant nucleosome as EM02, and collected the tomogram of the mixed specimen containing recombinant nucleosome core particle and nucleosomes with linker DNA as EM03. The detailed information of these datasets is summarized in Table S1.

### Detailed implementation in real nucleosome datasets

#### NMA of nucleosomes

For the NMA data processing of nucleosomes in this study, a series of atomic models were generated from nucleosomes with 147 bp DNA (PDB: 3AFA) by using a linear relationship between the amplitudes of normal modes 7 and 13. A gradual transition between the two ends, which represented a continuum of nucleosome conformations, was simulated. Equal random amplitudes uniformly distributed in the range [-250, 250] were used for the two normal modes 7 and 13. To visualize obvious continuous conformational changes, nucleosomes with long linker DNA were also studied. The atomic model of the 177 bp nucleosome was generated from PDB (7DBP) by removing the chain of H1. The following NMA was performed in a similar process as in the study of nucleosomes with 147 bp DNA.

#### Training model for prediction

To predict the tomograms of real nucleosome data, we used the simulated data that mimicked the real collection conditions to train the model. The SNR of the simulated data was also ensured to be close to the real data. Specifically, the training data were deposited into cubic subvolumes of 64 voxels at a pixel spacing of 4.44 Å. The training pairs were normalized before training. The other steps are described above.

#### Tomographic reconstruction

For the recombinant nucleosome datasets, the tilt movies were processed in Warp (Tegunov and Cramer, 2019), and the generated stacks were aligned using IMOD (Kremer et al., 1996). The reconstructions were generated from Warp using a pixel spacing of 4.44 Å, which was as same as that in the training dataset.

#### Predicting the real tomograms

The tomograms were normalized before prediction. After training, the model was used to predict the tomogram from Warp. The IsoNet prediction strategy of prediction was implemented. We split the entire tomogram into small subvolumes in 64 voxels to predict them separately. Then, output 3D chunks were combined to produce the final output.

### Single-particle analysis of the mixed nucleosomes

We collected and processed the mixed samples with the single-particle method. After 2D classification and 3D classification with Relion, we selected one class which had the feature of nucleosome with DNA. After refinement in Relion, a 3.7 Å map was obtained as the reference for REST-restored density.

### Template matching in simulated tomograms

Since the ground truth of the real coordinates in the simulated tomograms has been already known, the calculated coordinate can be obtained through template matching in Dynamo (Castaño-Díez et al., 2012). Thus, by subtracting the real coordinates from the calculated coordinates in the X-, Y- and Z-directions, the shift of the corresponding particle coordinates could also be determined. At the same time, each particle returned a CC value during the calculation, and further comparison was made between the raw data and the result using REST.

### The 3D visualization

IMOD was used to visualize the 2D slices, and UCSF Chimaera (Pettersen et al., 2004) and UCSF ChimeraX (Goddard et al., 2018) were used to visualize the 3D tomograms and subvolumes. Schematics were drawn using Adobe Illustrator.

## Data and code availability

The code is available at https://github.com/Zhang-hn1125/REST-beta.

## Acknowledgments and funding sources

This work was supported by grants from the Chinese Ministry of Science and Technology (2017YFA0504700, 2021YFA1300100, 2018YFE0203300), the National Natural Science Foundation of China (31730023, 31521002, 31600691), and the Chinese Academy of Sciences (CAS) (XDB37010100). All EM data were collected and processed at the Centre for Bio-imaging (CBI), Institute of Biophysics (IBP), Chinese Academy of Sciences (CAS). We would like to thank Jianguo Zhang, Xing Jia, Xiaojun Huang, Boling Zhu, for their technical help and support with electron microscopy.

## Author contributions

P.Z. and H.Z. initiated the project. H.Z. designed research and wrote the code; Y.L., L.W., K.S. and K.B. supported the nucleosome samples. H.Z. performed the cryo-EM sample preparation. H.Z. and Y.L. performed data collection. H.Z., Y.L. and D.L. analysed the data. H.Z., Y.L. and P.Z. wrote the paper.

**The authors declare no competing interest**.

## Supplementary Figures and Tables

**Fig. S1.**
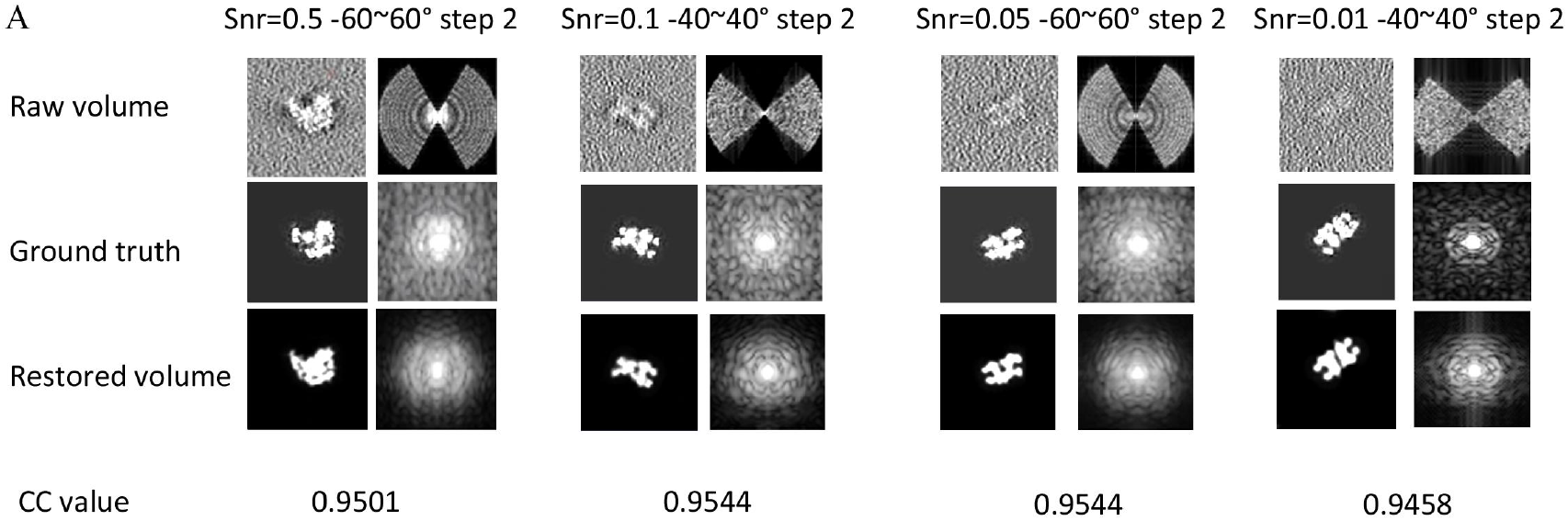
The performance of REST predictions under different simulation conditions. The difference in the data is shown in the first row. The left side of each pair is the XY-slice view, and the right side is the Fourier transform of the left particles. The particle represents the input, predict represents the results restored by REST and CC represents the cross coefficient between the predicted volume and ground truth.

**Fig. S2.**
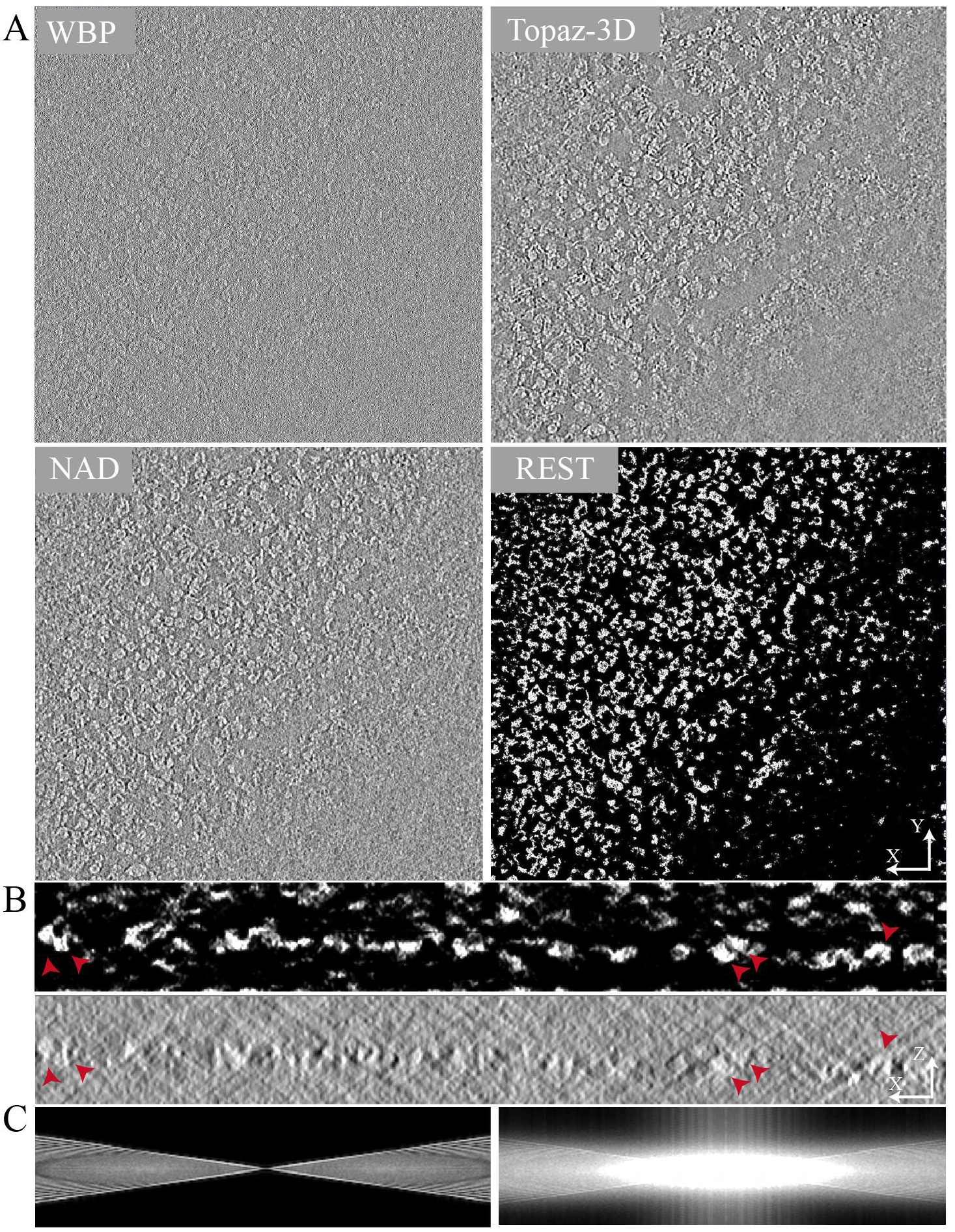
Enhancement of the visualization of nucleosomes in tomograms using REST. (A) Compared with the raw data, the NAD and Topaz-Denoise, REST methods can be used to thoroughly remove the noise shown in the tomographic slices. **(B)** The XZ-slice views of the REST-restored tomogram (top) and the denoised tomogram using Topaz-Denoise (bottom). **(C)** Fourier transforms of the raw tomogram (left) and REST-restored tomogram (right).

**Fig. S3.**
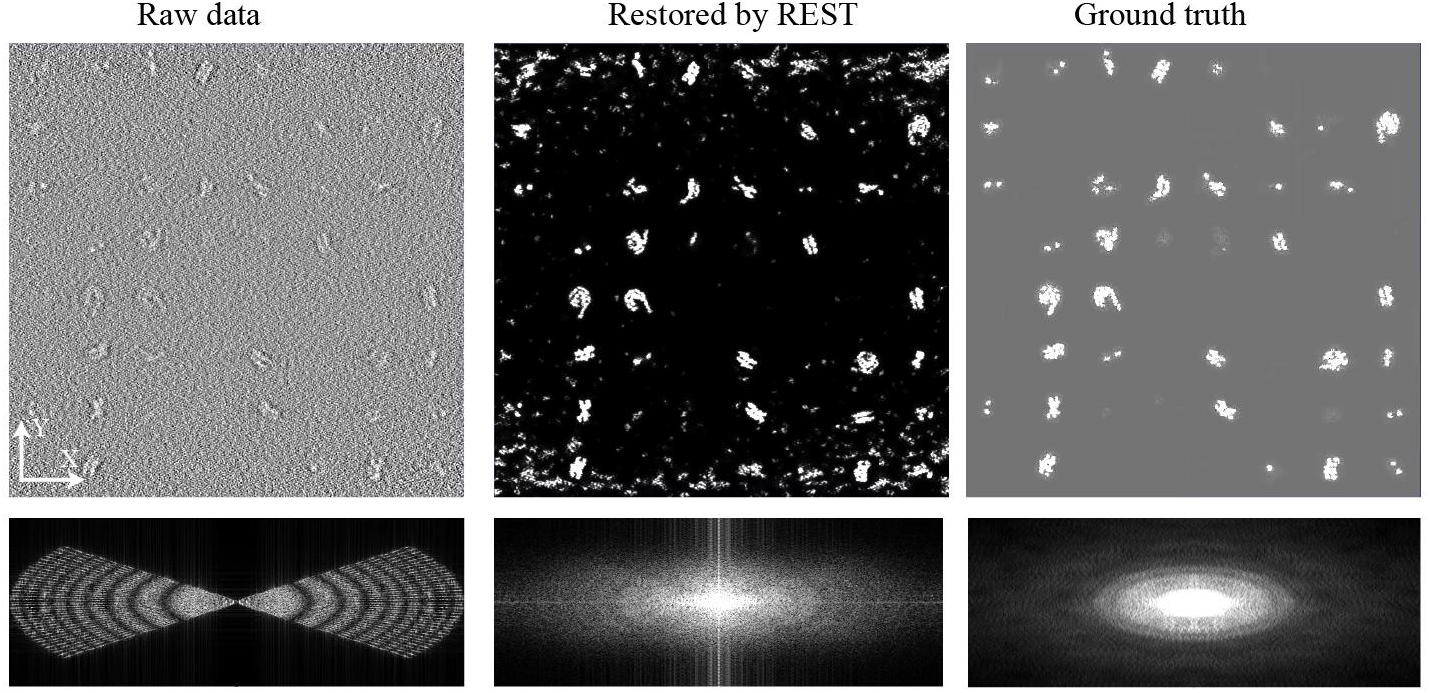
Top: The XY-slice views of the simulated raw tomogram (left), REST restored tomogram (middle) and the ground truth; Bottom: the corresponding Fourier transforms of the top tomograms.

**Fig. S4.**
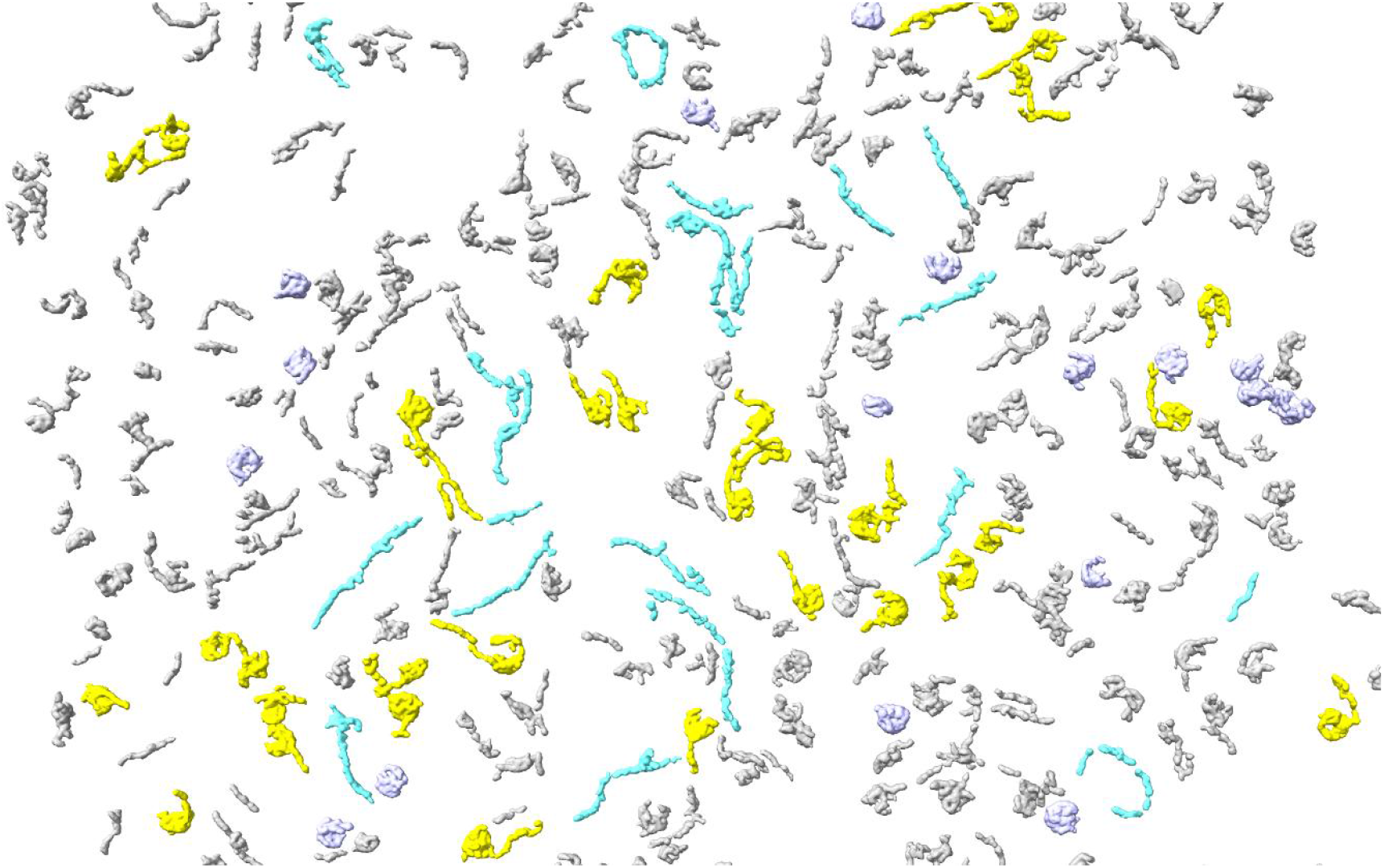
The whole tomogram of reconstituted nucleosomes restored by REST. A portion of nucleosome core particles, nucleosomes with linker DNA and free DNA are shown in purple, yellow and cyan, respectively.

**Table S1:**
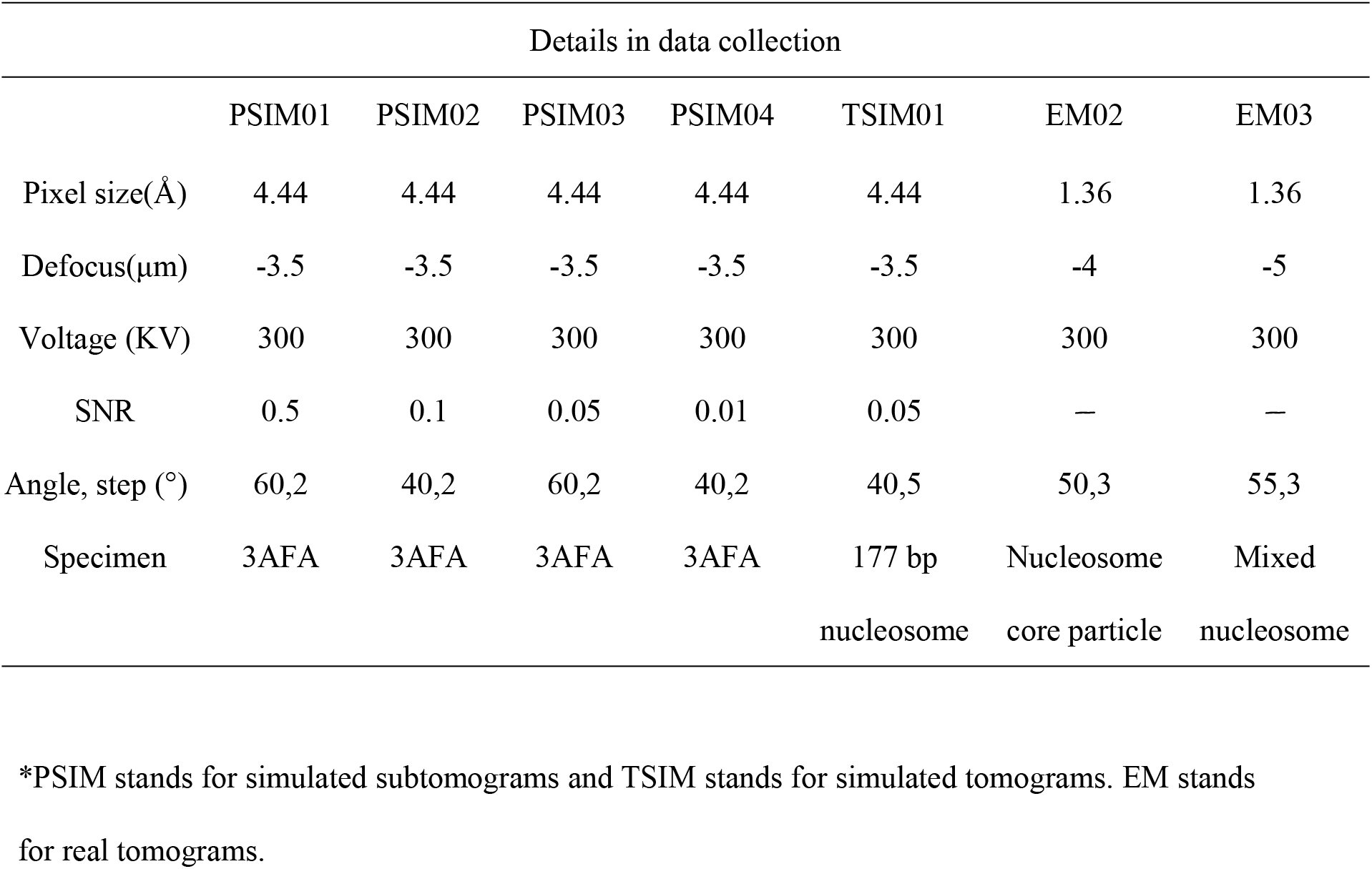
Detailed information about the simulated and real datasets.

**Table S2:** Cross coefficient (CC) between the T-I volume and restored volume shown in Fig. 6.

|  | Particle type |  |  |
| --- | --- | --- | --- |
|  | Nucleosome core particles | Nucleosomes including linker DNA | Free DNA |
| Particle 1 | 0.81 | 0.64 | 0.91 |
| Particle 2 | 0.84 | 0.89 | 0.85 |
| Particle 3 | 0.78 | 0.72 | 0.89 |
| Particle 4 | 0.77 | 0.76 | — |
| Particle 5 | 0.78 | 0.57 | — |
| Particle 6 | 0.70 | 0.72 | — |

## Notes

https://github.com/Zhang-hn1125/REST-beta

### Competing Interest Statement

The authors have declared no competing interest.

